# In vivo CRISPRi screen identified lncRNA portfolio crucial for cutaneous squamous cell carcinoma tumor growth

**DOI:** 10.1101/2024.10.16.618774

**Authors:** Gyuhyeon Kim, Zurab Siprashvili, Xue Yang, Jordan M. Meyers, Andrew Ji, Paul A. Khavari, Luca Ducoli

## Abstract

Cutaneous squamous cell carcinoma (cSCC) accounts for 20% of all skin cancer deaths globally, making it the second-highest subtype of skin cancer. The prevalence of cSCC in humans, as well as the poor capacity for an efficient prognosis, highlights the need to uncover alternative actors and mechanisms at the foundation of skin cancer development. Significant advances have been made to better understand some key factors in cSCC progression. However, little is known about the role of noncoding RNAs, particularly of a specific category called long noncoding RNA (lncRNA). By performing pseudobulk analysis of single-cell sequencing data from normal and cSCC human skin tissues, we determined a global portfolio of lncRNAs specifically expressed in keratinocyte subpopulations. Integration of CRISPR interference screens in vitro and the xenograft model identified several lncRNAs impacting the growth of cSCC cancer lines both in vitro and in vivo. Among these, we further validated LINC00704 and LINC01116 as proliferation-regulating lncRNAs in cSCC lines and potential biomarkers of cSCC progression. Taken together, our study provides a comprehensive signature of lncRNAs with roles in regulating cSCC progression.

## Introduction

The skin consists of three layers: epidermis, dermis, and subcutaneous tissue. The epidermis outer layer, primarily comprised of keratinocytes, is a protective barrier against physical damage, chemical stimulation, and microbial invasion. However, this barrier can be compromised by several factors, including chemicals, UV radiation, and pollutants, potentially leading to skin damage or cancer^1,2^. Skin cancer, a highly prevalent disease in humans, is typically classified into melanoma and non-melanoma categories, with the latter category including basal cell carcinoma (BCC) and cutaneous squamous cell carcinoma (cSCC)[PMID: 15660110]. cSCC is the second most prevalent skin cancer and accounts for 20% of skin cancer-related deaths worldwide^3–5^. cSCC’s aggressive nature makes it the leading cause of death among non-melanoma cancers, with advanced cases having a poor 10-year survival rate of less than 20%^6–8^. cSCC is generally believed to originate from keratinocytes, typically due to ultraviolet irradiation from the sunlight. Key risk factors include mutations in tumor suppressor genes like RAS, MYC, and p53^9^. While recent therapeutic advancements have ameliorated patient outcomes, the biology of cSCC progression is still an active area of study. In this context, great efforts have been invested in uncovering various regulatory mechanisms involved in the progression of cSCC at the signaling, epigenetic, transcriptional, and post-transcriptional levels that drive cSCC proliferation, survival, migration, and invasion^10–12^. However, limited studies were undertaken to elucidate the role of a regulatory molecule termed long noncoding RNA (lncRNA) in cSCC. The relationship between long non-coding RNA (lncRNA) expression and cancer metastasis has led to the concept that lncRNA profiles could serve as indicators for predicting cancer progression and patient outcomes^13^. Studies have shown increased expression of certain lncRNAs in ccSCC, including LINC00162 (also known as PICSAR), LINC00319, and THOR. Conversely, LINC00520, GAS5, and TINCR were shown to have reduced expression levels^14^. To expand our knowledge of the role of lncRNAs in the regulation of cSCC progression, we first reanalyzed previously single-cell sequencing datasets derived from normal and cSCC human tissues^15^. By performing pseudobulk analysis, we determined the landscape of lncRNAs specifically expressed in the three keratinocyte subpopulations (basal, cycling, differentiating keratinocytes) as well as on the recently uncovered cSCC-specific subpopulation (a.k.a TSK)^15^. CRISPR interference (CRISPRi) screen in cSCC cell lines as well as in xenograft cSCC model led to the identification of several lncRNAs influencing the growth of cSCC cell lines both in vitro and in vivo. Among the several candidates, we further validated the biological implication of LINC00704 and LINC01116 for their growth phenotype using antisense oligonucleotides, tissue microarray single-molecule RNA-FISH showed that these lncRNAs possess biomarker potential since their expression showed an cSCC-progression-dependent expression.

## Results

### cSCC keratinocyte subpopulations express a specific cohort of lncRNAs

To characterize lncRNAs in cSCC, we reanalyzed previously published human cSCC tumors and normal tissue single-cell RNA-seq (scRNA-seq) data (PMID:32579974). We first remapped the single-cell data using the latest lncRNA annotation from the FANTOM CAT database^16^. Secondly, we performed pseudobulk analysis by comparing within each other the three keratinocyte subpopulations (basal, cycling, and differentiating) present in normal and tumor samples and the tumor-specific keratinocytes (TSK) described in^15^ using the DESeq2 R package^17^ (**Fig. 1A and Supplementary Fig. 1A**). Principal component analysis (PCA) and Pearson correlation of the pseudobulk samples showed clear differences in the transcriptome between normal and tumor keratinocytes subpopulations as well as to TSK subpopulation (**Supplementary Fig. B-C**). Moreover, TSK markers were also expressed more strongly in the TSK pseudobulk samples compared to any other keratinocyte subpopulation either from normal or tumor samples (**Supplementary Fig. 1D**). We selected only lncRNAs that showed significantly higher expression in tumor-derived cSCC subpopulations against at least one keratinocyte subpopulation in both normal and tumor conditions (**Fig. 1B**). This analysis led to the identification of a specific cohort of 294 lncRNAs to be differentially expressed in the tumor cSCC subpopulations (**Fig. 1C, Supplementary Table 1**). Upset plot analysis showed that the vast majority of the identified lncRNAs are more highly expressed in the respective cSCC subpopulation (**Fig. 1D**), which aligns with previous knowledge that cells might display a specific set of lncRNAs that are important for their functions^18^. Overall, we found 116, 56, 39, and 30 lncRNAs to show generally higher expression in TSK, basal, cycling, and differentiating keratinocytes, respectively.

**Figure 1:**
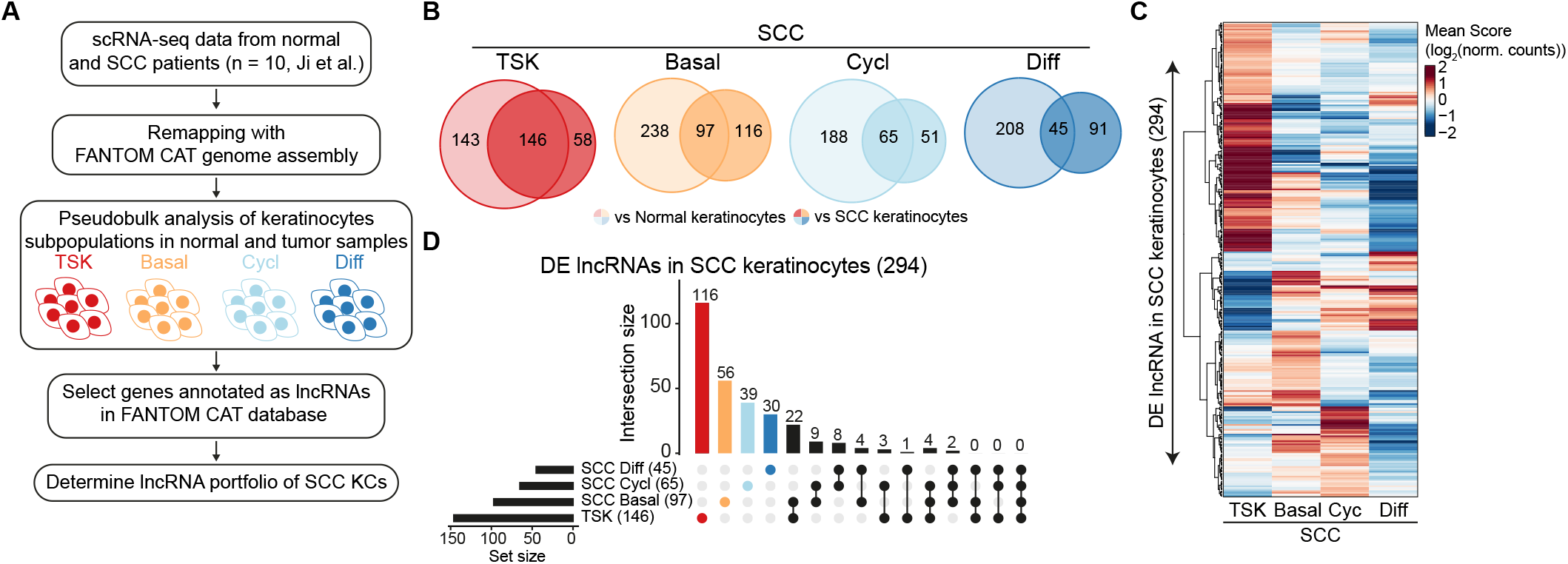
cSCC keratinocyte subpopulations express a specific cohort of lncRNAs. (**A**) Schematic representation of the pseudobulk analysis using DESeq2^17^ of human cSCC single-cell data from^15^. (**B**) Venn diagrams indicate the overlap of lncRNAs differentially expressed (DE) between the keratinocyte subpopulations in tumor and normal tissue. (**C**) Heatmap indicates the normalized pseudobulk counts of the 294 lncRNAs across the four cSCC keratinocyte subpopulations. Hierarchical clustering was performed using Euclidean distance and ward.D2 clustering algorithm (**D**) Upsetplot indicates the overlap of the 294 lncRNAs between the cSCC subpopulations.

### Selection of cSCC-associated lncRNAs displaying putative function

To identify putatively functional lncRNAs, we leveraged the functional feature annotation from the FANTOM CAT database^16^: “DNA conservation,” “GWAS trait associations,” “eQTL data,” and “Cell type ontology” (**Fig. 2A**). Of the 241 lncRNAs with higher expression in the respective cSCC keratinocyte subpopulations, we found 87 TSK, 37 basal, 31 cycling, and 21 differentiating to possess at least two functional features from the FANTOM CAT database^16^ (**Fig. 2B, Supplementary Table 2**). We, therefore, decided to focus on these 176 putative functional lncRNAs in our in vitro CRISPR screen experiments. Interestingly, the TSK-associated putative functional lncRNAs were particularly enriched for intergenic lncRNAs, a class of lncRNAs shown to have a higher probability of displaying functionality^16,19^ (**Fig. 2C**). Moreover, the TSK subpopulation showed the highest fraction of putative functional lncRNAs to show significantly higher expression against each of the cSCC keratinocyte subpopulations, not only suggesting that TSK display indeed a unique cohort of putative functional lncRNAs but also reinforcing their tumor-specific occurrence (**Fig. 2D**).

**Figure 2:**
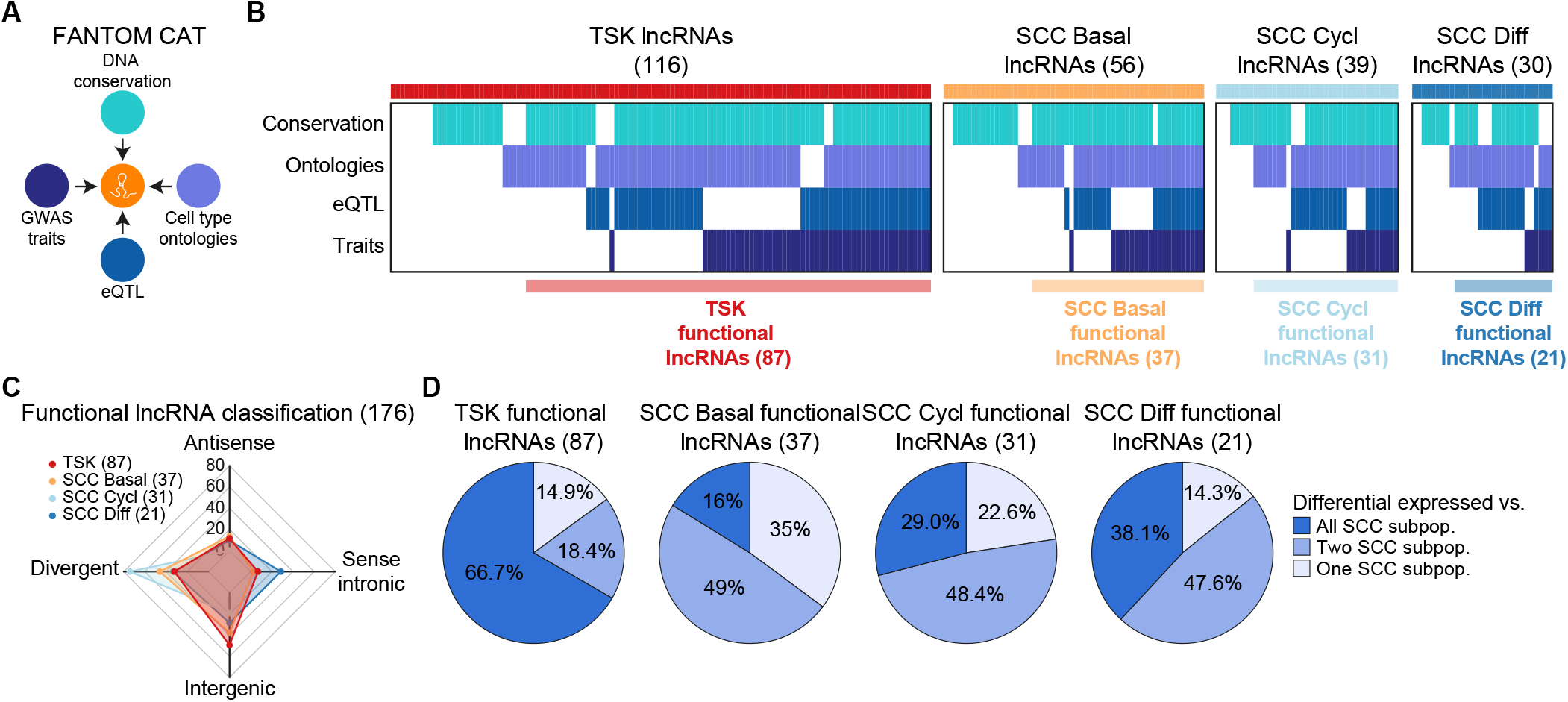
Selection of cSCC-associated lncRNAs displaying putative function. (**A**) Schematic representation of the criteria adopted to select functionally annotated lncRNAs in the FANTOM CAT database^16^. (**B**) Heatmap indicates the functional features of cSCC subpopulation-associated lncRNAs according to the FANTOM CAT database^16^. (**C**) Radarplot indicates the percentage of genomic categories (antisense, divergent, intergenic, and sense intronic) of cSCC subpopulation-associated lncRNAs. (**D**) Pie charts indicate the proportions of TSK, cSCC basal, cSCC cycling, and cSCC differentiating putative functional lncRNAs differentially expressed against other cSCC keratinocyte subpopulations.

### in vitro CRISPRi screen identifies candidates affecting cSCC proliferation in 2D

To investigate the biological functions of these lncRNA candidates in tumor growth on a high throughput scale, we used the CRISPR interference (CRISPRi) screen technology using dCas9-BFP-KRAB^20^ that showed the low off-target effect rate in targeting lncRNA transcripts^21,22^. This CRISPRi system showed the expected shift in western blot analysis and effectively knocked down epidermal differentiation markers in cSCC cell lines (**Supplementary Fig. 2A-B**). For the 176 cSCC putative functional lncRNAs, we designed 10 sgRNAs per target across a region from -200 and +450 nucleotides from each transcriptional starting site (TSS) of the cSCC lncRNAs annotated in the FANTOM CAT database^16^ (**Fig. 3A and Supplementary Fig. 2C**). In addition to safe targeting controls, we also included in our library several positive controls that were determined to influence cSCC tumor growth^15^. This brought a total of 9590 sgRNAs (**Supplementary Table 3**) cloned into two libraries (TSK library – TSK-related lncRNAs; KC library – basal, cycling, and differentiating lncRNA targets) (**Fig. 3B**). Analysis of TSK and KC library quality via miSeq showed that both libraries were of high quality with a narrow skew value of 2.34 and 2.1, respectively (**Supplementary Fig. 2D**).

**Figure 3:**
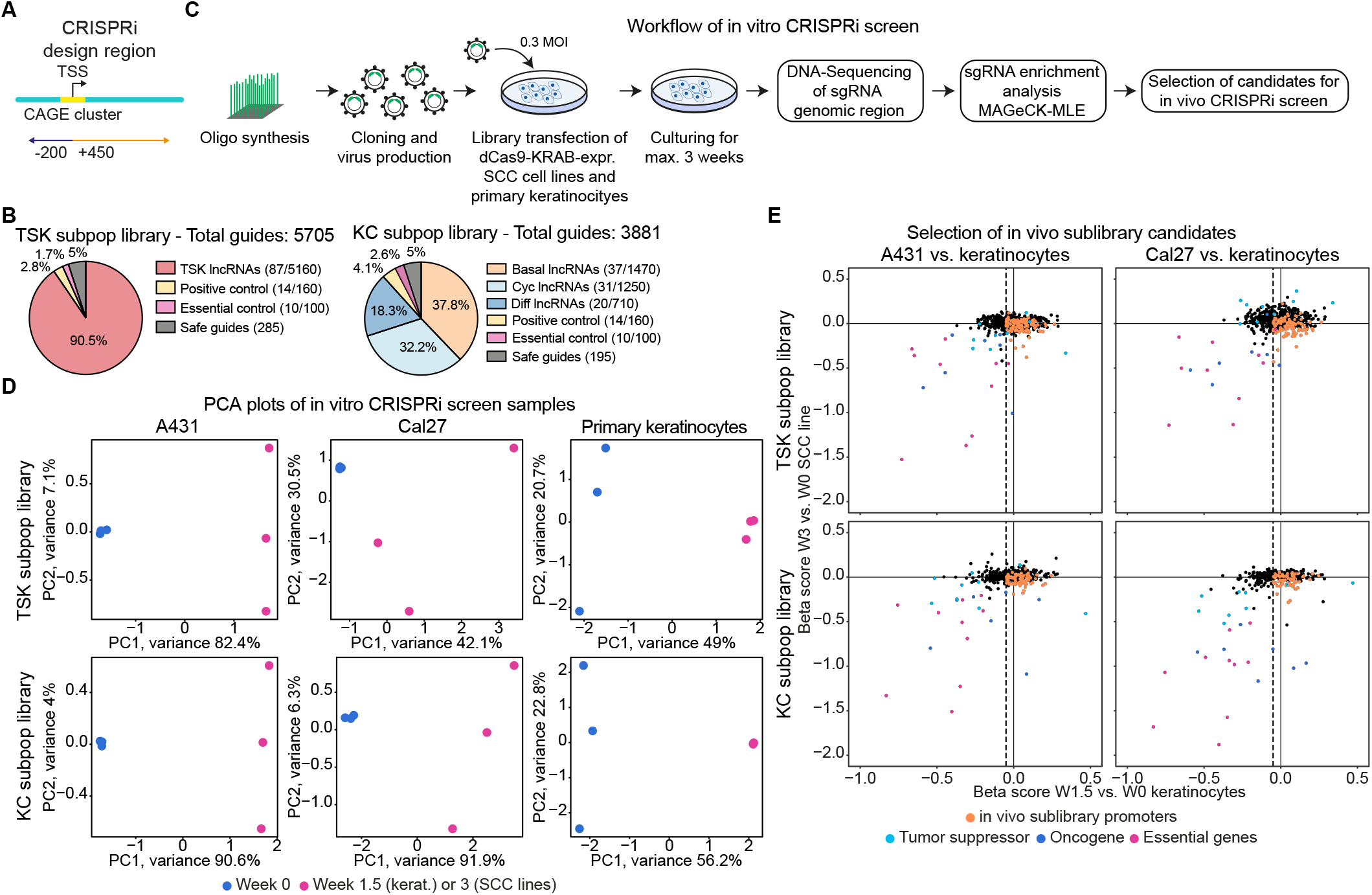
in vitro CRISPRi screen identifies candidates affecting cSCC proliferation in 2D. (**A**) Schematic representation of CRISPRi sgRNA design region around the transcriptional start site (TSS). (**B**) Pie charts indicate the composition of the TSK and KC subpopulation libraries. (**C**) Schematic representation of the in vitro CRISPRi screen experiment. (**D**) Principal component analysis (PCA) plots showing sample clustering. (**E**) Scatter plot of beta score values versus week 0, calculated using the MAGeCK-MLE algorithm^23^, showing the selection of relevant promoters to be included in the in vivo screen.

We then performed in vitro pooled CRISPRi knockdown followed by DNA sequencing in two cSCC lines, A431 and Cal27, as well as human primary keratinocytes (**Fig. 3C**). Using the MAGECK-MLE algorithm^23^, we then analyzed the impacts of lncRNA candidate knockdowns on the growth of the different cell types. We analyzed the changes in the samples of week 3 for cSCC cell lines and week 1.5 for primary keratinocytes against the respective day 0 cells. PCA analysis showed the expected separation between the samples (**Fig. 3D**). Most positive controls showed negative enrichment after the screen. Based on the enrichment analysis, we identified some discardable targets that showed negatively enriched in primary keratinocytes or positively enriched in cSCC lines. To do so, we selected lncRNA promoters with a β-score < -0.05 and a P-value < 0.1 in at least one cancer cell line but not in the primary keratinocytes (**Fig. 3E**). After additional filtering for robust expression in the scRNA-seq data^15^, we found a total of 176 lncRNA promoter targets (108 promoters from 39 lncRNAs from TSK library; 68 promoters from 28 lncRNAs from KC library) remains to be proceeded into in vivo CRISPRi screen (**Fig. 3E, Supplementary Table 4**).

### In vivo CRISPRi screens identify lncRNA affecting 3D cSCC tumor growth

To perform the in vivo CRISPRi screen, we first generated the library targeting the 176 lncRNA promoters (10 sgRNA per target promoter), termed in vivo sublibrary (**Fig. 4A and Supplementary Table 3**). The guide representation quality was checked by miSeq where we observed a narrow skew value of 1.74 (**Supplementary Fig. 3A**). Then, we injected three mice per cSCC cell line (A431 and Cal27) with cells previously infected with the CRISPRi system and the in vivo library. After 3-4 weeks, we harvested the tumors and performed DNA isolation. Isolated DNA was then subjected to library preparation and DNA sequencing (**Fig. 4B, Supplementary Fig. 3B)**. The correlation between tumor weight and volume, as well as DNA amount and tumor weight, was observed (**Supplementary Fig. 3C-D)**. The expected separation between the samples was observed after PCA analysis (**Fig. 4C**).

**Figure 4:**
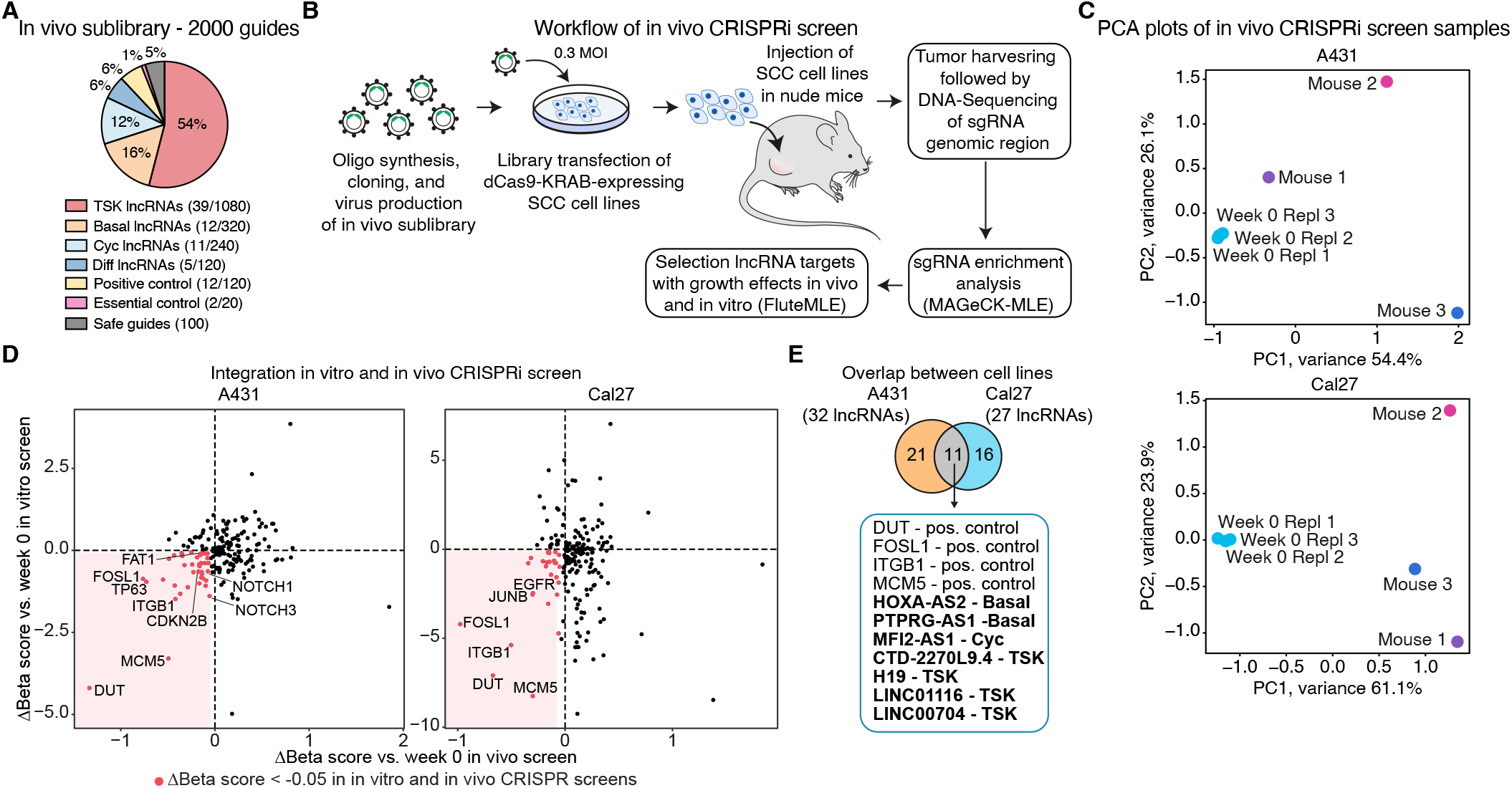
In vivo CRISPRi screens identify lncRNAs affecting 3D cSCC tumor growth. (**A**) Pie chart indicate the composition of the in vivo library. (**B**) Schematic representation of the in vivo CRISPRi screen experiment. (**C**) PCA plots showing sample clustering. (**D**) Scatter plots indicate normalized Δbeta score values versus week 0, calculated using the MAGeCK-MLE followed by FluteMLE^24^, in vivo and in vitro for A431 (left) and Cal27 (right). Red dots indicate targets with Δbeta score values < -0.05 in both in vitro and in vivo data. (**E**) The Venn diagram indicates the overlap between A431 and Cal27 CRISPRi screens selected lncRNAs and positive controls.

To determine the growth impact of lncRNA targets, we first calculated the sgRNA enrichment ß-score using the MAGeCK-MLE algorithm^23^. Using the FluteMLE R package^24^, we integrated the in vitro and in vivo CRISPR screen results by calculating the difference between normalized ß-scores of week 0 and tumor/week 3 samples (**Fig. 4D**). As expected, many positive control knockouts showed a reduction in the growth of cSCC cell lines both in vivo and in vitro (**Fig. 4D**). We found 31 and 22 lncRNAs to show a congruent effect on growth in vitro and in vivo for A431 and Cal27 (**Supplementary Table 5**). Overlap between cSCC lines lncRNAs found 7 of them to consistently affect the growth of both A431 and Cal27 in vivo as well as in vitro (**Fig. 4F**). Remarkably, some of these lncRNAs, LINC00704 and LINC01116 (AC017048.3), have been previously associated with different types of cancers^25,26^ (**Figure 3**).

### Validate the role of the TSK-signature lncRNA LINC00704 in cSCC progression

Four of seven lncRNAs are explicitly expressed in the TSK subpopulation among four cSCC subpopulations, suggesting them as potential cell-type signatures. To test this, we reanalyzed previously published single-cell datasets from the Cal27 xenograft model^15^. After remapping with the FANTOM CAT annotation^16^, we performed a correlation analysis to reveal whether the putative oncogenic lncRNAs showed a significant co-expression with previously defined markers for each of the four cSCC subpopulations. Among the 7 lncRNA hits, we found that LINC00704 and LINC01116 significantly correlated with a large fraction of markers for the TSK subpopulations (**Fig. 5A**). Dimension reduction analysis of Cal27 xenograft model showed that LINC00704 and LINC01116 expression were indeed colocalizing with TSK known makers, suggesting that those lncRNAs might represent a specific functional signature of TSK subpopulations (**Fig. 5B**). Similar analysis in single-cell data of transformed-keratinocytes spheroids overexpressing CDK4(R24C) and Ras(G12V) constitutive mutations showed that LINC00704 was also co-expressed with TSK markers in this settings (**Supplementary Fig. 4A-B**).

**Figure 5:**
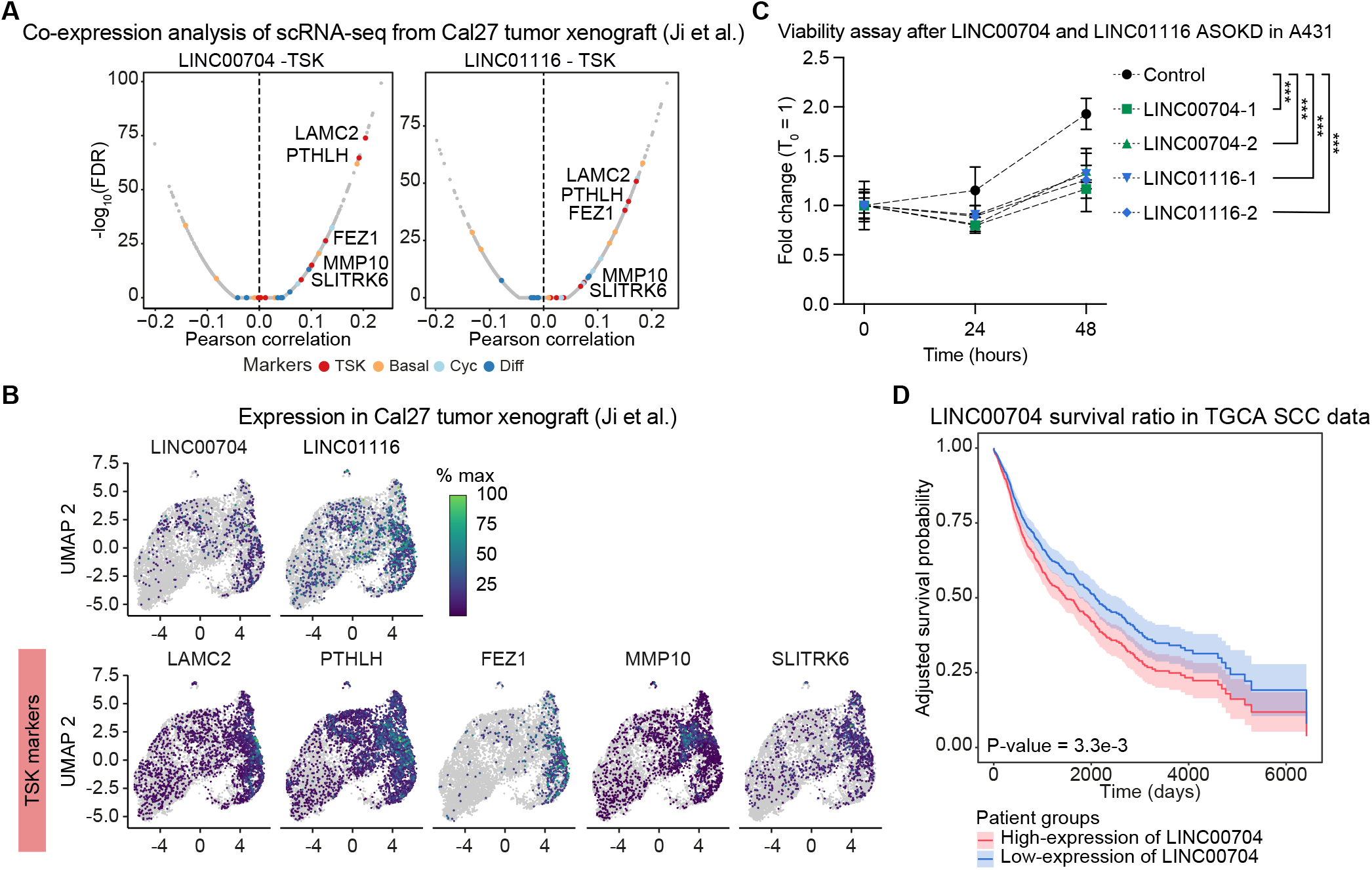
Validate the role of the TSK-signature lncRNA LINC00704 in cSCC progression. (**A**) Pearson correlation analysis indicates the co-expression analysis results of LINC00704 and LINC01116 with TSK subpopulation markers in Cal27 xenograft single-cell sequencing data^15^. (**B**) UMAP plots indicate the expression distribution in Cal27 cells isolated from xenograft tumors of LINC00704 and LINC01116 and TSK markers. (**C**) Line plot indicates the fold change against time point 0 of A431 viability measured by cell titer blue after the knockdown via ASO of LINC00704 and LINC01116. Data are presented as mean values + SD (n=6). Two-way ANOVA with Dunnett’s multiple comparisons test; ***, P-value < 0.001. (**D**) Kaplan-Meyer (MK) plot indicates the adjusted survival probability for gender bias of cSCC patients retrieved from TCGA expressing high (red) or low (blue) levels of LINC00704. 95% confidence intervals for survival time are shown.

To validate the involvement of those candidates in cSCC cell proliferation, we first designed two antisense oligonucleotides (ASOs) targeting the mature transcripts of LINC00704 and LINC01116 (**Supplementary Fig. 4C**). qPCR analysis after ASO-transfection confirmed the knockdown efficiency for both ASOs (**Supplementary Fig. 4D**). Time course cell-titer blue viability assay after the ASO-mediated knockdown of both LINC00704 and LINC01116 in A431 cells validated the involvement of LINC00704 and LINC01116 in the regulation of cSCC line viability (**Fig. 5C**).

To understand the extent to which the expression of LINC00704 and LINC01116 are affecting cSCC progression, we performed a TGCA survival analysis using data generated from squamous cell carcinoma tissues. We found that, after correcting for gender, high expression of LINC00704 displayed a significant reduction in patient survival (**Fig. 5D**). In the case of LINC01116, a potential trend is observed but with limited statistical significance. Overall, these analyses supports LINC00704 to be a robust biomarker in cSCC progression.

## Discussion

Collectively, our results provide new insights into the involvement of lncRNAs in cSCC progression. Through pseudobulk analysis using single-cell sequencing data, we curated the list of lncRNAs explicitly expressed in cSCC keratinocyte subpopulations. Then, we identified the set of lncRNAs affecting cSCC growth by in vitro and in vivo CRISPR screens and further validated LINC00704 as an exemplary putative carcinogenic target. Using CRISPRi screens both in vitro and in a mouse xenograft model provides robust validation of our findings, addressing potential limitations of in vitro studies alone. This approach allows for identifying lncRNAs that are functionally important in a more physiologically relevant context, considering the complex 3D tumor microenvironment.

Identifying lncRNAs with subpopulation-specific expression patterns underscores the heterogeneity within cSCC tumors, a characteristic that has been increasingly recognized as crucial in cancer biology^27^. In fact, our results suggest that different keratinocyte subpopulations within cSCC tumors may have distinct lncRNA profiles. This reinforces the general idea that lncRNAs are molecular signatures of cellular states^28^. In particular, the tumor-specific keratinocyte (TSK) subpopulation displayed the highest number of explicitly expressed lncRNAs. This supports the notion that this population is a unique cancer cell population that does not recapitulate normal epidermal states, unlike other subpopulations found in cSCC^15,29,30^. TSK is typically localized to a fibrovascular niche at the leading edges of the tumor, suggesting their involvement in invasion and metastasis. TSKs serve as a hub for intercellular communication within the tumor microenvironment, engaging in extensive ligand-receptor interactions with surrounding stromal and immune cells^15^. Recent studies have shown that in recurrent cSCC, TSKs exhibit significant epithelial-mesenchymal transition (EMT) features, which may be attributed to their active communication with cancer-associated fibroblasts^29^. The presence and characteristics of TSKs underscore the complexity of tumor heterogeneity in cSCC biology. Identifying lncRNAs specifically expressed in this cell population affecting tumor growth may provide a mechanistic insight into subpopulation-specific contribution to tumor growth and enable future therapeutic approaches targeting specific keratinocyte subpopulations through lncRNAs.

Among the lncRNAs explicitly expressed in the subpopulation, we identified seven candidates that show impairment of cSCC growth both in vitro and in vivo. Four of them were specifically expressed in the TSK subpopulation. Two of them, LINC00704 and LINC01116, showed robust co-expression with TSK markers in xenograft and spheroids models (for LINC00704) and significant influence on cell viability. These results are consistent with previous studies that have implicated both lncRNAs in various cancer types, including nasopharyngeal carcinoma and lung adenocarcinoma^26,31^. Our study suggests its implication on cSCC growth, specifically through cSCC cancer-specific subpopulations.

The observed impact on cell viability aligns with their reported functions in other cancers, further supporting their role as a key player in cancer progression across different tumor types. LINC00704, also known as MANCR (Mitotically Associated Long Non-coding RNA), has been previously associated with cell cycle regulation and genomic stability in nasopharyngeal carcinoma (NPC)^26^. Here, overexpression of LINC00704 promoted NPC cell proliferation, migration, and invasion^26^. In papillary thyroid carcinoma (PTC), LINC00704 was also found to be elevated and functioned as an oncogene by promoting cell proliferation, migration, and invasion^32^. LINC01116, on the other hand, has been reported to regulate critical cellular processes such as the epithelial-mesenchymal transition (EMT) and cell cycle progression in lung adenocarcinoma^31^. Although not in line with our TCGA analysis in cSCC tumors, high expression of LINC01116 was associated in this cancer type with poor prognosis, suggesting a cancer context dependency^31^. In glioma, LINC01116 has been shown to promote tumor growth and neutrophil recruitment by regulating IL-1β expression^33^.

Analysis of TCGA data revealed significantly reduced survival for cSCC patients with high LINC00704 expression, further emphasizing its potential clinical relevance in cSCC. In fact, LINC00704 could serve as a biomarker for cSCC progression or for distinguishing between different keratinocyte subpopulations within tumors. The potential of lncRNAs as cancer biomarkers has been demonstrated in various cancer types^34^, and our findings extend this possibility to cSCC. In conclusion, these findings increase our understanding of lncRNA involvement in cSCC biology, particularly in the context of tumor heterogeneity and growth. Identifying subpopulation-specific lncRNAs with essential functions in tumor growth offers new avenues for targeted therapies that could disrupt tumor progression while minimizing effects on normal tissue. Thus, future studies should focus on understanding the extent to which these lncRNAs contribute to the tumorigenic potential of the TSK subpopulation in cSCC progression, with the aim to improve diagnosis, prognosis, and treatment selection, and, ultimately, patient outcomes.

## Acknowledgments

We thank M. Pilo and A. Dazey for expert administrative assistance, and members of the Khavari lab for helpful discussions. This work was supported by NIAMS/NIH grants AR045192 and AR076965 to P.A.K. and by USVA Merit Review grant BX001409 to P.A.K., and by Swiss National Science Foundation Postdoc Mobility Fellowship P500BP-203019 to L.D.

## Author Contributions

G.K., P.A.K., and L.D. conceived the project. G.K., Z.S., X.Y, and L.D. performed experiments. G.K. and L.D. performed data analysis. A.J. supported data interpretation. P.A.K. and L.D. guided experiments and data analysis. G.K., P.A.K., and L.D. wrote the manuscript with input from all co-authors.

## References

(1) Poirier, M. C. Chemical-Induced DNA Damage and Human Cancer Risk. Discov. Med. 2012, 14 (77), 283–288.

(2) Luch, A. Nature and Nurture – Lessons from Chemical Carcinogenesis. Nat. Rev. Cancer 2005, 5 (2), 113–125.

(3) Leiter, U.; Keim, U.; Garbe, C. Epidemiology of Skin Cancer: Update 2019. In Sunlight, Vitamin D and Skin Cancer; Reichrath, J., Ed.; Springer International Publishing: Cham, 2020; pp 123–139.

(4) Lomas, A.; Leonardi-Bee, J.; Bath-Hextall, F. A Systematic Review of Worldwide Incidence of Nonmelanoma Skin Cancer. Br. J. Dermatol. 2012, 166 (5), 1069–1080.

(5) Rogers, H. W.; Weinstock, M. A.; Harris, A. R.; Hinckley, M. R.; Feldman, S. R.; Fleischer, A. B.; Coldiron, B. M. Incidence Estimate of Nonmelanoma Skin Cancer in the United States, 2006. Arch. Dermatol. 2010, 146 (3), 283–287.

(6) Pollock, J. Cutaneous Squamous-Cell Carcinoma. N. Engl. J. Med. 2001, 345 (4), 296; author reply 296-297. PMID: 11474682.

(7) Cassarino, D. S.; DeRienzo, D. P.; Barr, R. J. Cutaneous Squamous Cell Carcinoma: A Comprehensive Clinicopathologic Classification. J. Cutan. Pathol. 2006, 33 (4), 261–279.

(8) Hillen, U.; Ulrich, M.; Alter, M.; Becker, J. C.; Gutzmer, R.; Leiter, U.; Lonsdorf, A.; Messerschmidt, A.; Ulrich, C. Kutanes Plattenepithelkarzinom unter Berücksichtigung besonderer Patientengruppen. Hautarzt 2014, 65 (7), 590– 599.

(9) Thompson, A. K.; Kelley, B. F.; Prokop, L. J.; Murad, M. H.; Baum, C. L. Risk Factors for Cutaneous Squamous Cell Carcinoma Recurrence, Metastasis, and Disease-Specific Death: A Systematic Review and Meta-Analysis. JAMA Dermatol. 2016, 152 (4), 419–428.

(10) Cozma, E.-C.; Banciu, L. M.; Soare, C.; Cretoiu, S.-M. Update on the Molecular Pathology of Cutaneous Squamous Cell Carcinoma. Int. J. Mol. Sci. 2023, 24 (7), 6646.

(11) Jiang, R.; Fritz, M.; Que, S. K. T. Cutaneous Squamous Cell Carcinoma: An Updated Review. Cancers 2024, 16 (10), 1800.

(12) Khavari, P. A. Modelling Cancer in Human Skin Tissue. Nat. Rev. Cancer 2006, 6 (4), 270–280.

(13) Frontiers | Long Non-coding RNAs in Cancer: Implications for Diagnosis, Prognosis, and Therapy. https://www.frontiersin.org/journals/medicine/articles/10.3389/fmed.2020.61239 3/full (accessed 2024-10-15).

(14) Wang, Y.; Sun, B.; Wen, X.; Hao, D.; Du, D.; He, G.; Jiang, X. The Roles of lncRNA in Cutaneous Squamous Cell Carcinoma. Front. Oncol. 2020, 10.

(15) Ji, A. L.; Rubin, A. J.; Thrane, K.; Jiang, S.; Reynolds, D. L.; Meyers, R. M.; Guo, M. G.; George, B. M.; Mollbrink, A.; Bergenstråhle, J.; Larsson, L.; Bai, Y.; Zhu, B.; Bhaduri, A.; Meyers, J. M.; Rovira-Clavé, X.; Hollmig, S. T.; Aasi, S. Z.; Nolan, G. P.; Lundeberg, J.; Khavari, P. A. Multimodal Analysis of Composition and Spatial Architecture in Human Squamous Cell Carcinoma. Cell 2020, 182 (2), 497-514.e22.

(16) Hon, C.-C.; Ramilowski, J. A.; Harshbarger, J.; Bertin, N.; Rackham, O. J. L.; Gough, J.; Denisenko, E.; Schmeier, S.; Poulsen, T. M.; Severin, J.; Lizio, M.; Kawaji, H.; Kasukawa, T.; Itoh, M.; Burroughs, A. M.; Noma, S.; Djebali, S.; Alam, T.; Medvedeva, Y. A.; Testa, A. C.; Lipovich, L.; Yip, C.-W.; Abugessaisa, I.; Mendez, M.; Hasegawa, A.; Tang, D.; Lassmann, T.; Heutink, P.; Babina, M.; Wells, C. A.; Kojima, S.; Nakamura, Y.; Suzuki, H.; Daub, C. O.; de Hoon, M. J. L.; Arner, E.; Hayashizaki, Y.; Carninci, P.; Forrest, A. R. R. An Atlas of Human Long Non-Coding RNAs with Accurate 5′ Ends. Nature 2017, 543 (7644), 199– 204.

(17) Love, M. I.; Huber, W.; Anders, S. Moderated Estimation of Fold Change and Dispersion for RNA-Seq Data with DESeq2. Genome Biol. 2014, 15 (12), 550.

(18) Ward, M.; McEwan, C.; Mills, J. D.; Janitz, M. Conservation and Tissue-Specific Transcription Patterns of Long Noncoding RNAs. J. Hum. Transcr. 2015, 1 (1), 2–9. PMID: 27335896.

(19) Rodriguez-Lopez, M.; Anver, S.; Cotobal, C.; Kamrad, S.; Malecki, M.; Correia-Melo, C.; Hoti, M.; Townsend, S.; Marguerat, S.; Pong, S. K.; Wu, M. Y.; Montemayor, L.; Howell, M.; Ralser, M.; Bähler, J. Functional Profiling of Long Intergenic Non-Coding RNAs in Fission Yeast. eLife 2022, 11, e76000.

(20) Tian, R.; Gachechiladze, M. A.; Ludwig, C. H.; Laurie, M. T.; Hong, J. Y.; Nathaniel, D.; Prabhu, A. V.; Fernandopulle, M. S.; Patel, R.; Abshari, M.; Ward, M. E.; Kampmann, M. CRISPR Interference-Based Platform for Multimodal Genetic Screens in Human iPSC-Derived Neurons. Neuron 2019, 104 (2), 239-255.e12.

(21) Stojic, L.; Lun, A. T. L.; Mangei, J.; Mascalchi, P.; Quarantotti, V.; Barr, A. R.; Bakal, C.; Marioni, J. C.; Gergely, F.; Odom, D. T. Specificity of RNAi, LNA and CRISPRi as Loss-of-Function Methods in Transcriptional Analysis. Nucleic Acids Res. 2018, 46 (12), 5950–5966.

(22) Smith, I.; Greenside, P. G.; Natoli, T.; Lahr, D. L.; Wadden, D.; Tirosh, I.; Narayan, R.; Root, D. E.; Golub, T. R.; Subramanian, A.; Doench, J. G. Evaluation of RNAi and CRISPR Technologies by Large-Scale Gene Expression Profiling in the Connectivity Map. PLOS Biol. 2017, 15 (11), e2003213.

(23) Li, W.; Köster, J.; Xu, H.; Chen, C.-H.; Xiao, T.; Liu, J. S.; Brown, M.; Liu, X. S. Quality Control, Modeling, and Visualization of CRISPR Screens with MAGeCK-VISPR. Genome Biol. 2015, 16 (1), 281.

(24) Wang, B.; Wang, M.; Zhang, W.; Xiao, T.; Chen, C.-H.; Wu, A.; Wu, F.; Traugh, N.; Wang, X.; Li, Z.; Mei, S.; Cui, Y.; Shi, S.; Lipp, J. J.; Hinterndorfer, M.; Zuber, J.; Brown, M.; Li, W.; Liu, X. S. Integrative Analysis of Pooled CRISPR Genetic Screens Using MAGeCKFlute. Nat. Protoc. 2019, 14 (3), 756–780.

(25) Xu, Y.; Yu, X.; Zhang, M.; Zheng, Q.; Sun, Z.; He, Y.; Guo, W. Promising Advances in LINC01116 Related to Cancer. Front. Cell Dev. Biol. 2021, 9.

(26) Wang, Y.; Tan, Q.-Y.; Shen, Y.; Liu, C.-Y.; Huang, T.; Huai, D.; Dai, J. LINC00704 Contributes to the Proliferation and Accelerates the Cell Cycle of Nasopharyngeal Carcinoma Cells via Regulating ETS1/CDK6 Axis. Kaohsiung J. Med. Sci. 2022, 38 (4), 312–320.

(27) Cyll, K.; Ersvær, E.; Vlatkovic, L.; Pradhan, M.; Kildal, W.; Avranden Kjær, M.; Kleppe, A.; Hveem, T. S.; Carlsen, B.; Gill, S.; Löffeler, S.; Haug, E. S.; Wæhre, H.; Sooriakumaran, P.; Danielsen, H. E. Tumour Heterogeneity Poses a Significant Challenge to Cancer Biomarker Research. Br. J. Cancer 2017, 117 (3), 367–375.

(28) Johnsson, P.; Ziegenhain, C.; Hartmanis, L.; Hendriks, G.-J.; Hagemann-Jensen, M.; Reinius, B.; Sandberg, R. Transcriptional Kinetics and Molecular Functions of Long Noncoding RNAs. Nat. Genet. 2022, 54 (3), 306–317.

(29) Li, X.; Zhao, S.; Bian, X.; Zhang, L.; Lu, L.; Pei, S.; Dong, L.; Shi, W.; Huang, L.; Zhang, X.; Chen, M.; Chen, X.; Yin, M. Signatures of EMT, Immunosuppression, and Inflammation in Primary and Recurrent Human Cutaneous Squamous Cell Carcinoma at Single-Cell Resolution. Theranostics 2022, 12 (17), 7532–7549.

(30) Amôr, N. G.; Santos, P. S. da S.; Campanelli, A. P. The Tumor Microenvironment in SCC: Mechanisms and Therapeutic Opportunities. Front. Cell Dev. Biol. 2021, 9.

(31) Zeng, L.; Lyu, X.; Yuan, J.; Wang, W.; Zhao, N.; Liu, B.; Sun, R.; Meng, X.; Yang, S. Long Non-Coding RNA LINC01116 Is Overexpressed in Lung Adenocarcinoma and Promotes Tumor Proliferation and Metastasis. Am. J. Transl. Res. 2020, 12 (8), 4302–4313. PMID: 32913506.

(32) Lin, Y.; Jiang, J. Long Non-Coding RNA LINC00704 Promotes Cell Proliferation, Migration, and Invasion in Papillary Thyroid Carcinoma via miR-204-5p/HMGB1 Axis. Open Life Sci. 2020, 15 (1), 561–571. PMID: 33817244.

(33) Wang, T.; Cao, L.; Dong, X.; Wu, F.; De, W.; Huang, L.; Wan, Q. LINC01116 Promotes Tumor Proliferation and Neutrophil Recruitment via DDX5-Mediated Regulation of IL-1β in Glioma Cell. Cell Death Dis. 2020, 11 (5), 1–13.

(34) Badowski, C.; He, B.; Garmire, L. X. Blood-Derived lncRNAs as Biomarkers for Cancer Diagnosis: The Good, the Bad and the Beauty. Npj Precis. Oncol. 2022, 6 (1), 1–18.

