## Supplementary Information for "In vivo CRISPRi screen identified lncRNA portfolio crucial for cutaneous squamous cell carcinoma tumor growth"

#### Supplementary figure legends

**Supplementary Figure 1: Pseudobulk analysis of scRNA-seq data from human skin tissue samples.**

(**A**) Schematic representation of the tests performed by DESeq2[1] in the pseudobulk analysis. (**B**) Heatmap indicates the pearson correlation values calculated from normalized psuedobulk counts derived from the keratinocytes subpopulation identified in the 10 normal and cSCC tissue samples from[2]. Hierarchical clustering was performed using Euclidean distance and ward.D2 clustering algorithm. (**C**) Principal component analysis of normalized pseudobulk counts for each keratinocyte subpopulation of the 10 normal and cSCC tissue samples from[2]. (**D**) Barplot indicates the $log_2$FC values of tumor-specific keratinocytes (TSK) markers in TSK compared to the keratinocyte subpopulation basal, cyclin, and differentiation.

**Supplementary Figure 2: Generation of CRISPRi library to target cSCC lncRNAs**

(**A**) Western blot images indicate the FLAG and Actin signal in keratinocytes infected with normal Cas9 and dCas9-BFP-KRAB. (**B**) Bar plot indicates the fold change against safe targeting control of several keratinocyte-related gene expressions in A431 and Cal27 cells. (**C**) Box plots indicate the number of transcripts (left) and transcriptional start sites (TSS, right) in the different cSCC keratinocyte subpopulations. (**D**) Line plots indicate the cumulative fraction distribution of sgRNAs in the TSK and KC subpopulation libraries. Skew values represent the ratio between the 90% and 10% sgRNA feature counts.

**Supplementary Figure 3: Generation of in vivo library and mouse xenograft experiment**

(**A**) Line plots indicate the cumulative fraction distribution of sgRNAs in “in vivo” libraries. Skew values represent the ratio between the 90% and 10% sgRNA feature counts. (**B**) Images indicate the location and size of the tumor xenografts in nude mice at harvesting time (3.5 weeks for A431 and 4.5 weeks for Cal27). The pictures of tumors were taken after being arranged approximately by their volumes upon the harvests. (from top left to bottom right). The tumor lable is associated with the top panel and weight of each tumor is provided (**C**) Scatter plot indicates the tumor volume and weight of the six tumors isolated from three nude mice from each SCC cell lines. (**D**) The scatter plot indicates the amount of harvested DNA and tumor weight of the six tumors isolated from three mice from each SCC cell line.

**Supplementary Figure 4: LINC00704 co-expression analysis in spheroids, ASO efficiency, and LINC01116 TCGA analysis.**

(**A**) Pearson correlation analysis indicates the co-expression analysis results of LINC00704 with TSK subpopulation markers in transformed keratinocytes spheroids overexpressing CDK4(R24C) and Ras(G12V) mutations. (**B**) UMAP plots indicate the expression distribution in transformed keratinocyte spheroids of LINC00704 and LINC01116 and TSK markers. (**C**) Schematic representation of ASO location (orange lines) across the mature transcript of LINC00704 and LINC01116. (**D**) Barplots indicate the fold change in the expression of LINC00704 and LINC01116 after their knockdown using two ASOs (n=2). (**E**) Kaplan-Meyer (MK) plot indicates the adjusted survival probability for gender bias of cSCC patients retrieved from TCGA expressing high (red) or low (blue) levels of LINC01116. 95% confidence intervals for survival time are shown.

#### Suppl. Figure 1: Pseudobulk analysis of scRNA-seq data from human skin tissue samples

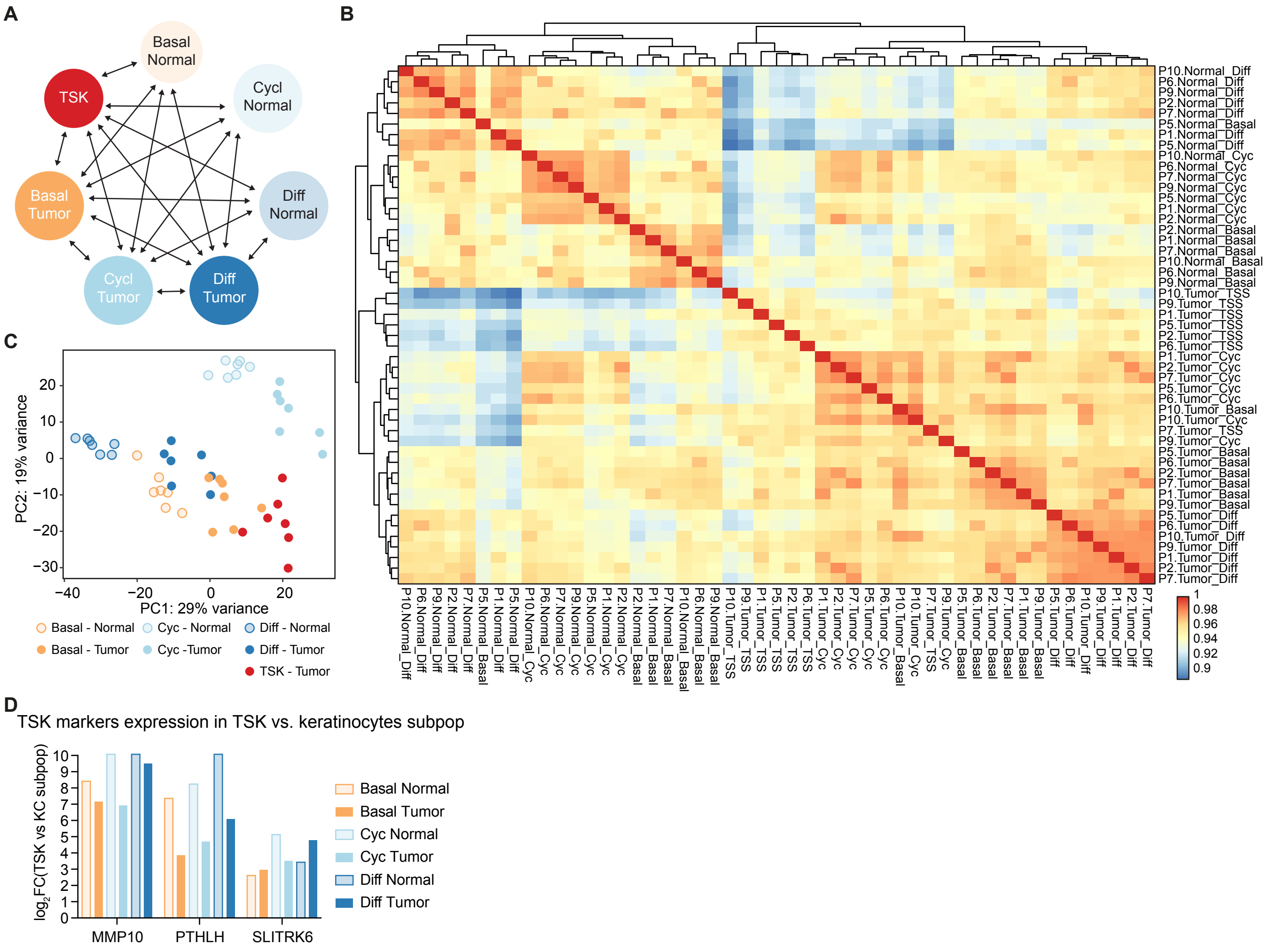

### Suppl. Figure 2: Generation of CRISPRi library to target SCC lncRNAs.

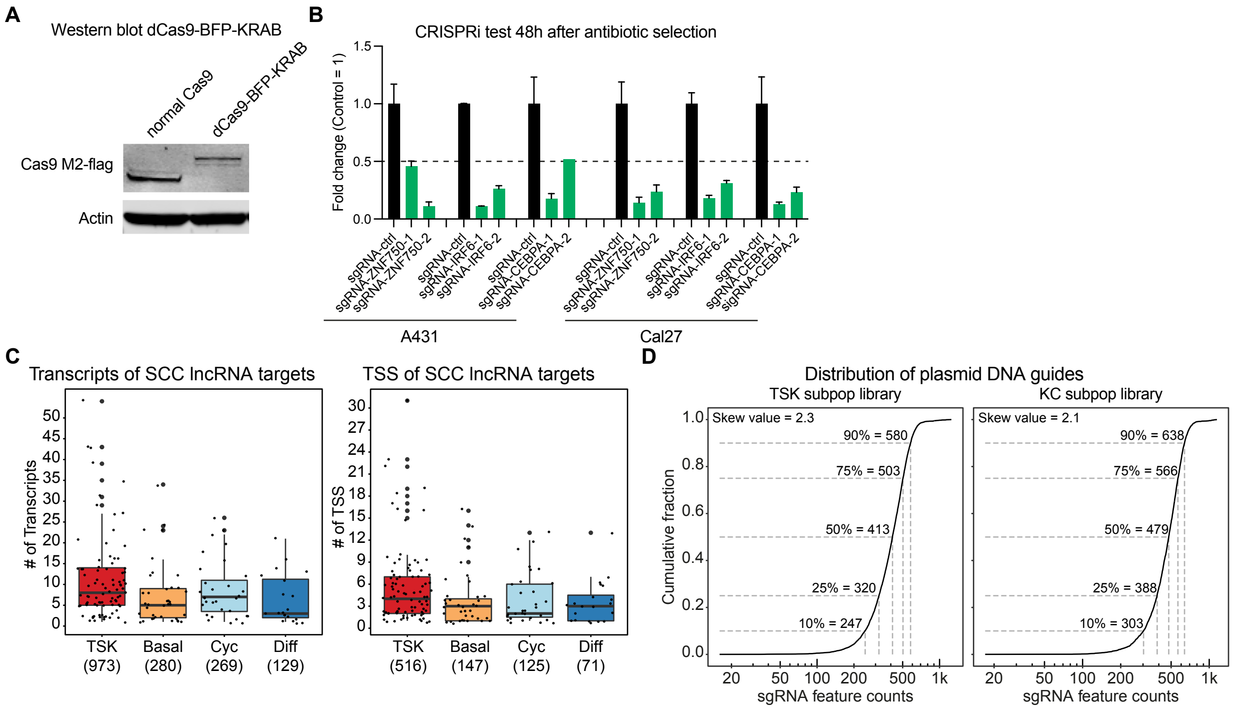

### Suppl. Figure 3: Generation of in vivo library and mouse xenograft experiment.

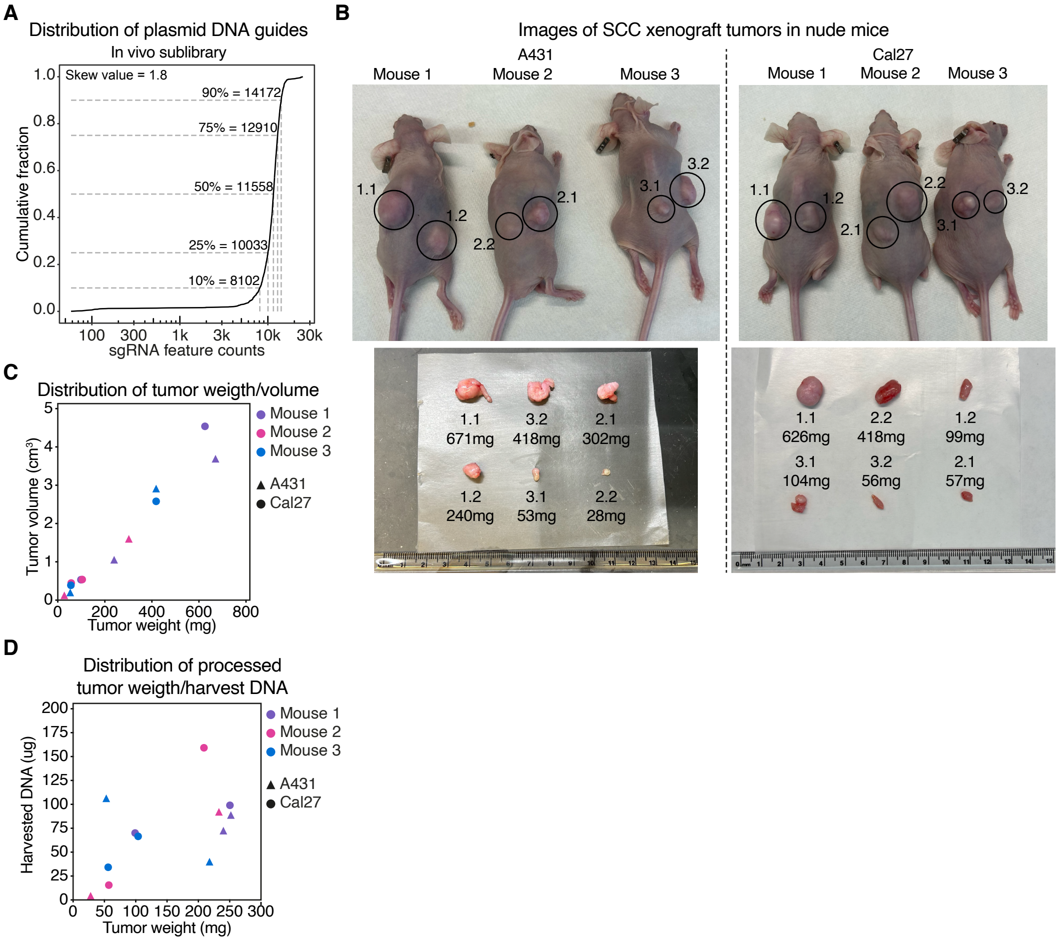

**Suppl. Figure 4: LINC00704 co-expression analysis in spheroids, ASO efficiency, and LINC01116 TCGA analysis.**

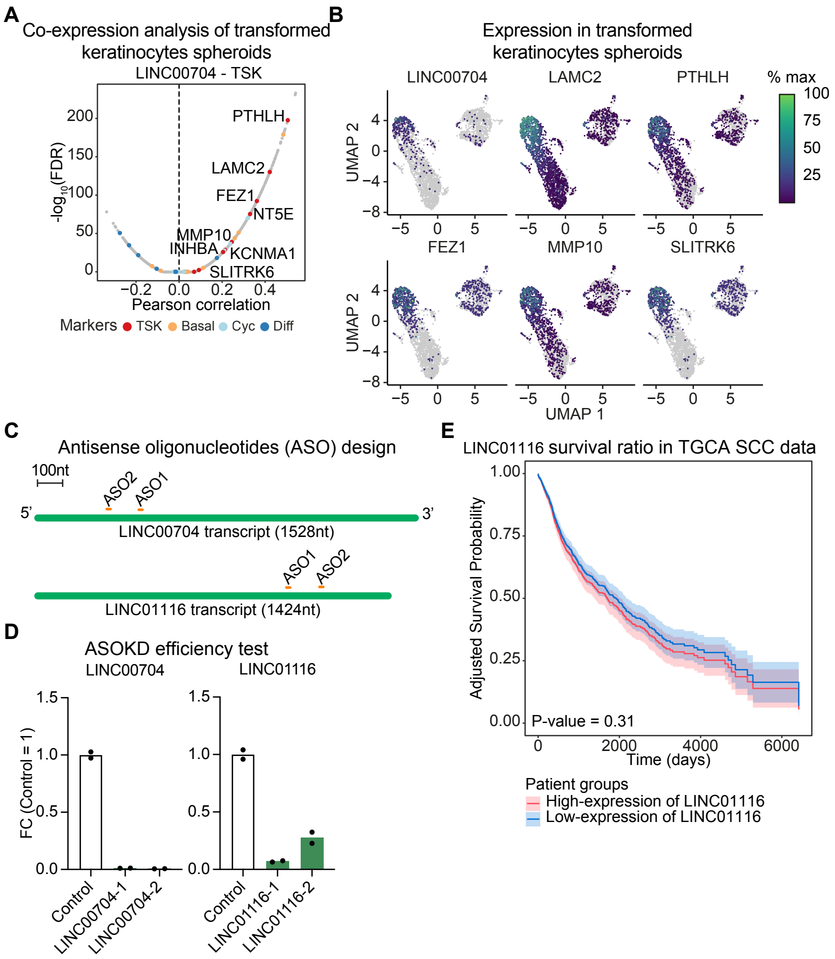

#### Methods

##### Cell culture

The human A431 and Cal27 cell lines were cultured as described in the previous study[2]. Primary human keratinocytes were obtained from fresh, surgically discarded neonatal foreskin, cultured in Keratinocyte-SFM and Medium 154 with Keratinocyte supplement, and subcultured regularly before reaching high confluency for a few cycles.

##### RNA isolation, reverse transcription, and qPCR

To perform quantitative RT-PCR (qRT-PCR), total RNA was extracted using RNeasy Plus mini kit (QIAGEN), and concentration was quantified using NanoDrop ONE (Thermo Fisher). According to the manufacturer's instructions, equal amounts of total RNA (usually 1µg) were reverse transcribed using the iScript cDNA Synthesis Kit (Bio-Rad). qRT-PCR analysis was performed using the LightCycler 480 II System (Roche) with the SYBR Green Master Mix (Thermo Fisher). Samples were run in triplicates (10ng cDNA per reaction) and normalized to levels of GAPDH for each reaction. qPCR Primers are listed in **Supplementary Table 6**.

##### Western blotting

For immunoblot analysis, sample protein concentrations were determined using the Pierce BCA Protein Assay kit (ThermoFisher) following the manufacturer’s instructions. 20 µg of cell lysates were loaded per lane on a NUPAGE 4-12% Bis-Tris SDS-PAGE gel (ThermoFisher) and transferred to a pre-activated PVDF membrane (Promega) at 4°C. The membrane was blocked with Intercept blocking buffer (PBS, LICOR Biosciences) at room temperature for 1 hour. The membrane was then incubated with rabbit anti-flag antibody (CST) at a dilution of 1:1000 at 4°C overnight. The membrane was then washed five times for 5min in each time with TBS-T and incubated with secondary goat anti-rabbit antibodies (LI-COR Biosciences) at a dilution of 1:5,000 for 1 hour at room temperature. After incubation with secondary antibodies, membranes were washed again three times for 5min in each time with TBS-T and once with DPBS and visualized using the Odyssey CLx Infrared Imaging System (LI-COR Biosciences). Image processing was performed using Licor ImageStudioLite software (LI-COR Biosciences). Antibodies are listed in **Supplementary Table 6**.

##### Pseudobulk analysis of human skin tissue single-cell RNA-seq data

10x fastq data from[2] were aligned with `cellranger` using the transcriptome assembly downloaded from the FANTOM CAT database[3]. After importing the expression values using the Seurat R package[4], Pseudobulk expression matrices were built for each cell population derived from individual donors determined in[2] and subsequently normalized using the `estimateSizeFactors()` function from the DESeq2 package[1]. Differential expression analysis was performed using the `DESeq()` function. Differentially expressed lncRNAs between the normal and tumor SCC cell populations were defined as FDR < 0.01 and $\log_2$ fold change (FC) > 1. Only genes annotated as lncRNAs and with at least two functional features in the FANTOM CAT database[3] were kept for further analysis.

**In-vitro CRISPRi screen**

*Guide library construction.* We constructed two pooled oligo libraries and included ten guides per target and positive controls. transcriptional start sites, as annotated in the FANTOM CAT database[3], showing a TPM expression value higher than 25% quantile in the scRNA-seq data from normal and tumor skin tissues. The guides targeting these TSSs were designed using CRISPick[5]. The following libraries were generated: (1) KC library contains lncRNA targets selected from basal, cycling, and differentiating subpopulations. The library contains a total of 3885 guides. 3430 guides for 343 lncRNA targets, 260 guides for 26 positive controls, and 195 Safe targeting guides. (2) The TSK library contains lncRNA targets selected exclusively from the TSK subpopulation. The library contains a total of 5705 guides. 5160 guides for 516 lncRNA targets, 260 guides for 26 positive controls, and 285 Safe targeting guides.

*Guide library cloning*. A 69nt guide oligo pool was ordered from IDT with the following sequences. 5'- ATCTTGTGGAAAGGACGAAACACCG-[ guide spacer sequence] – GTTTAAGAGCTAAGCTGGAAACAG-3'. 10pmol of the lyophilized pooled oligo was resuspended into 100nM, and 4ul (8.6ng) of the oligo pool was then assembled with 118ng Esp3I digested pLentiguide vector using NEBuilder HiFi DNA Assembly. The reaction was transformed into Stellar competent cells and grown overnight in liquid culture with ampicillin selection. Following plasmid preparation using the Plasmid Plus Maxi Kit (QIAGEN), the library was sequenced on a MiSeq (Illumina, Inc.) to confirm guide representation.

*In-vitro CRISPRi screen*. Lentivirus has been produced in Lenti-X cells seeded higher than 70% in 10cm (or 15cm) dish. Cells in each 10cm dish were transfected with 7.5ug CMV, 3.5ug MDG, 7.5ug Plasmid in 625ul OptiMEM mixed with another tube containing 25ul

Lipofectamine in 625ul OptiMEM. (The volume or amount were doubled for cells in 15cm dish). The media was replaced after the overnight incubation, collected after every 24h with media replacement for two times, filtered through a 0.45um PES membrane, and concentrated with Lenti-X concentrator (Takara). Cas9 transfection was performed on A431, CAL27, and Primary Keratinocytes and selected with blasticidin. Pooled guide lentivirus was titrated in each of dCas9-KRAB expressing cell lines to achieve an MOI of 0.3. Following selection with puromycin for two to three days, the transfection rates were checked with Cell Titer Blue, and triplicate of cells (with +1000x coverage) were collected, flash frozen, and stored at -80C for each replicate as the initial screen time point. Cells were cultured by maintaining at least 1000x coverage with regular basis of subculturing for three weeks (Primary Keratinocyte for 1.5 week due to inherently limited proliferation capacity). DNA was isolated from the cell pellet using QIAGEN DNeasy Blood and Tissue Kit and the sequencing library was made as described in[2].

**In-vivo CRISPRi screen**

*Guide library design and cloning*. Based on the MaGeCK analysis performed on *in vitro* CRISPRi, promoters were annotated as hits if it showed $\beta < -0.05$ with p-value < 0.1 at least in one cancer cell line but not in primary keratinocyte. First, lncRNAs that don't contain any hit promoters were removed. Subsequently, promoters negatively enriched ($\beta < -0.05$, p value < 0.1) at primary keratinocytes or positively enriched ($\beta > 0.05$, p value < 0.1) at cancer cells were removed. For a particular set of seven lncRNAs (CASC15, LINC00152, LINC00704, LINC00707, RP11-85M11.2, RP11-701P16.5, PR1-272L16.1) mentioned in the publication with relevant contexts, we still included them in the final list by only excluding promoters negatively enriched ($\beta < -0.05$, p value < 0.1) at primary keratinocytes. Finally, promoters showing less than 5 TPM were removed, which resulted in a total of 176 promoter targets. (108 promoters from 39 lncRNAs from TSK library. 68 promoters from 28 lncRNAs from KC library). The same 10 guides targeting the 176 promoters for the 107 lncRNAs from the in vitro screen for each target were included. Together with 100 safe targeting guides and 140 control guides for 14 control genes, the final in vivo library resulted in total 2000 guides.

*In-vivo CRISPRi screen*. For the in vivo CRISPR screen, pooled oligo Lenti virus was made and transfected on dCas9-KRAB expressing A431 and CAL27 cell lines as described above. After the 3 days of selection in puromycin, four million cells in 100ul were gently mixed with 50ul Matrigel (Corning) until it turned into a homogenous mixture and injected with 28G insulin syringe into both sides of rear flanks of SCID Hairless Outbred (SHO®) mice (Charles River). Triplicate of cells (with +1000x coverage) were collected, flash frozen, and stored at -80C for each replicate as the initial screen time point. Three mice (a total of six tumors) were performed on each cell line. Based on the visual observation of subcutaneous tumor growths, the mice injected with the same cell lines were sacrificed on the same days after 3-4 weeks. (A431: 3 weeks, CAL27: 4 weeks) The tumors were surgically harvested, minced, and homogenized in an MP Biomedical lysing matrix A tube. Genomic DNA was isolated using QIAGEN DNeasy Blood and Tissue Kit, and the sequencing library was made as described in[6] with an adjustment of using <10g of gDNA in PCR1 reaction and a pair of primers matching with our sequences and manually determining PCR2 cycles based on a trial run to prevent overamplification.

**CRISPRi screen analysis**

In each screen, UMI was compressed using `umi_tools extract`, and adaptor sequences were removed using `cutadapt`. Subsequently, the remaining reads were aligned on the guide sequences using `bowtie` (maximum error: 1, minimum length: 16). The count for each guide was collected using `umi_tools count`, then transformed into scc format. For in vitro screen analysis, MaGeCK MLE was performed based on the guide counts with the design matrix of comparing the timepoint of the initial collection and the post-screen collection as described[7]. In vivo and in vitro CRISPR screens were integrated by first determining the ß scores of each sample against plasmid DNA determined by miSeq (Illumina) using the MaGeCK MLE algorithm[7]. ß scores were normalized and visualized using the MaGeCKFlute R package[8] following the authors' guidelines.

**Lentivirus production and keratinocyte infection**

To produce viruses, 5x$10^6$ HEK293T cells passaged in a 10 cm plate and cultured overnight. On the following day, HEK293T cells were transfected with 8µg of p8.91, 2µg pMDG, and 10µg pLEX-CDK4(R24C)-Ras(G12V)[9] with 44µL of p3000 reagent in 1mL Opti-MEM (Gibco) containing 55µL Lipofectamine 3000 (ThermoFisher). The media was replaced after 16 hours. Viral-containing media was collected 48 hours after transfection, filtered through a 0.45 um PES membrane, and concentrated to 50X with the Lenti-X concentrator (Takara).

For infection, the virus titer was first determined using the Lenti-X Go-Stix Plus kit (Takara), according to the manufacturer's instructions. 1x$10^6$ Primary human keratinocytes were infected with an MOI of 1.0 in DMEM (Gibco) containing 10% FBS with polybrene (5 µg/mL) and incubated overnight. The next day, fresh media was added, and cells were selected using puromycin (1 µg/mL, Gibco) for 72 hours before performing downstream experiments.

**Single-cell RNA-seq of transformed keratinocytes spheroids**

A 10mg/mL solution of Poly(2-hydroxyethyl methacrylate) (pHEMA, sigma) was prepared in 95% ethanol and warmed at 37°C until fully dissolved. Two layers of PHEMA were applied to 10cm dishes by spreading 5mL of the solution and allowing it to dry under a tissue culture hood for 24 hours per layer. The coated dishes were sterilized under UV light for at least 15 minutes prior to use. For spheroid formation, CDK4(R24C)-Ras(G12V) cells were seeded into pHEMA-coated 10cm dishes at a final concentration of $2.5x10^5$ cells/mL in KGM medium for spheroid formation. The cultures were incubated at 37°C in a 5% CO2 incubator for a week. Spheroids were then collected in a 15mL falcon tube and reduced to single-cell suspension using 0.25% trypsin with DNase I (0.2 mg/mL, Worthington) and Collagenase I (5 mg/mL, Gibco) for 30 minutes at 37°C. Cells were washed 3 times in DMEM containing 10% FBS. Cells were collected by centrifugation at 450g for 5min and finally resuspended in PBS (Gibco) containing 0.04% BSA. Cell count was determined using a Countess machine (Thermo Fisher). After passing through a cell strainer (Corning), 25’000 cells were loaded into a 10x Chromium machine (10x genomics). Library preparation was performed using the Chromium GEM-X Single Cell 3’ Kit (10x Genomics), followed by sequencing at Novogene.

**Co-expression analysis from single-cell sequencing data**

Single-cell RNA-seq data from Cal27 xenograft[2] and CDK4(R24C)-Ras(G12V) transformed spheroids were mapped as described above. Seurat objects were generated, and expression values were normalized using the SCTransform() function from the Seurat R package[4]. Correlation between lncRNA candidates and each gene across the profiled cells was performed using the normalized expression values and the cor.test() R function. The generated p-values were corrected using the p.adjust() R function.

**ASO design and efficiency test**

Two locked nucleic acids (LNA) phosphorothioate GapmeRs per lncRNA target were designed using the MyDesign online tool (Qiagen) using the transcripts ENST00000430998 (LINC00704) and ENST00000295549 (LINC01116). To test the knockdown efficiency of each ASO, we performed a reverse transfection using Lipofectamine RNAiMAX (Thermo Fisher Scientific). In 10cm dishes, 2mL Opti-MEM,

20nM of scrambled control ASO or two ASOs targeting LINC00704 and LINC01116, and 25µL Lipofectamine RNAiMAX (Thermo) were mixed and incubated for 20min. Then, $1x10^6$ A431 cells per dish were seeded into each 10cm dish and cultured for 48h in a 5% $CO_2$ incubator. After 48h, cells were lysed in LRT plus lysis buffer (Qiagen), and RNA isolation was performed using an RNeasy mini kit (Qiagen) following the manufacturer's instructions. Knockdown efficiency for each ASO was checked by qPCR, as described above, using the primers listed in **Supplementary Table 6**. GAPDH was used as the housekeeping gene.

**CellTiter-blue (CTB) viability assay**

$1 \times 10^6$ A431 cells were reverse transfected with Lipofectamine RNAi MAX as described above and cultured for 48h in a 5% $CO_2$ incubator. After 48h, transfected A431 cells were detached and seeded at a 15000 cells/well density into 96-well plate (Corning). At each time point, media was replaced with 100 µL CellTiter-blue (Promega) diluted 1:5 in complete media were added to each well. The plate was incubated for 45min at 37 °C. Finally, fluorescence intensities were measured using a SpectraMax M5 (Molecular Devices) and the SoftMax Pro 7 (ver. 7.1.0) using the cell-titer blue in-built protocol.

**TCGA survival analysis**

TCGA analysis was performed using the TCGAbiolinks R package[10]. Clinical information and expression data were downloaded for the following TCGA projects: CESC, ESCA, BLCA, LUSC, and HNSC using the `GDCquery_clinic()` and `GDCdownload()` functions. Expression data were normalized using DEseq2[1] using the `vst()` function. Cox proportional hazards regression model fitting corrected for gender bias was performed using the survival and adjustedCurves R packages.

#### Reference supplementary information

(1) Love, M. I.; Huber, W.; Anders, S. Moderated Estimation of Fold Change and Dispersion for RNA-Seq Data with DESeq2. *Genome Biol.* **2014**, *15* (12), 550.

(2) Ji, A. L.; Rubin, A. J.; Thrane, K.; Jiang, S.; Reynolds, D. L.; Meyers, R. M.; Guo, M. G.; George, B. M.; Mollbrink, A.; Bergenstråhle, J.; Larsson, L.; Bai, Y.; Zhu, B.; Bhaduri, A.; Meyers, J. M.; Rovira-Clavé, X.; Hollmig, S. T.; Aasi, S. Z.; Nolan, G. P.; Lundeberg, J.; Khavari, P. A. Multimodal Analysis of Composition and Spatial Architecture in Human Squamous Cell Carcinoma. *Cell* **2020**, *182* (2), 497-514.e22.

(3) Hon, C.-C.; Ramilowski, J. A.; Harshbarger, J.; Bertin, N.; Rackham, O. J. L.; Gough, J.; Denisenko, E.; Schmeier, S.; Poulsen, T. M.; Severin, J.; Lizio, M.; Kawaji, H.; Kasukawa, T.; Itoh, M.; Burroughs, A. M.; Noma, S.; Djebali, S.; Alam, T.; Medvedeva, Y. A.; Testa, A. C.; Lipovich, L.; Yip, C.-W.; Abugessaisa, I.; Mendez, M.; Hasegawa, A.; Tang, D.; Lassmann, T.; Heutink, P.; Babina, M.; Wells, C. A.; Kojima, S.; Nakamura, Y.; Suzuki, H.; Daub, C. O.; de Hoon, M. J. L.; Arner, E.; Hayashizaki, Y.; Carninci, P.; Forrest, A. R. R. An Atlas of Human Long Non-Coding RNAs with Accurate 5′ Ends. *Nature* **2017**, *543* (7644), 199–204.

(4) Satija, R.; Farrell, J. A.; Gennert, D.; Schier, A. F.; Regev, A. Spatial Reconstruction of Single-Cell Gene Expression Data. *Nat. Biotechnol.* **2015**, *33* (5), 495–502.

(5) Doench, J. G.; Fusi, N.; Sullender, M.; Hegde, M.; Vaimberg, E. W.; Donovan, K. F.; Smith, I.; Tothova, Z.; Wilen, C.; Orchard, R.; Virgin, H. W.; Listgarten, J.; Root, D. E. Optimized sgRNA Design to Maximize Activity and Minimize Off-Target Effects of CRISPR-Cas9. *Nat. Biotechnol.* **2016**, *34* (2), 184–191.

(6) Mathiowetz, A. J.; Roberts, M. A.; Morgens, D. W.; Olzmann, J. A.; Li, Z. Protocol for Performing Pooled CRISPR-Cas9 Loss-of-Function Screens. *STAR Protoc.* **2023**, *4* (2), 102201.

(7) Li, W.; Köster, J.; Xu, H.; Chen, C.-H.; Xiao, T.; Liu, J. S.; Brown, M.; Liu, X. S. Quality Control, Modeling, and Visualization of CRISPR Screens with MAGeCK-VISPR. *Genome Biol.* **2015**, *16* (1), 281.

(8) Wang, B.; Wang, M.; Zhang, W.; Xiao, T.; Chen, C.-H.; Wu, A.; Wu, F.; Traugh, N.; Wang, X.; Li, Z.; Mei, S.; Cui, Y.; Shi, S.; Lipp, J. J.; Hinterndorfer, M.; Zuber, J.; Brown, M.; Li, W.; Liu, X. S. Integrative Analysis of Pooled CRISPR Genetic Screens Using MAGeCKFlute. *Nat. Protoc.* **2019**, *14* (3), 756–780. PMID: 30710114.

(9) Lazarov, M.; Kubo, Y.; Cai, T.; Dajee, M.; Tarutani, M.; Lin, Q.; Fang, M.; Tao, S.; Green, C. L.; Khavari, P. A. CDK4 Coexpression with Ras Generates Malignant Human Epidermal Tumorigenesis. *Nat. Med.* **2002**, *8* (10), 1105–1114.

(10) Colaprico, A.; Silva, T. C.; Olsen, C.; Garofano, L.; Cava, C.; Garolini, D.; Sabedot, T. S.; Malta, T. M.; Pagnotta, S. M.; Castiglioni, I.; Ceccarelli, M.; Bontempi, G.; Noushmehr, H. TCGAbiolinks: An R/Bioconductor Package for Integrative Analysis of TCGA Data. *Nucleic Acids Res.* **2016**, *44* (8), e71.
